## Supplementary Figures and Tables for "LIVE-PAINT: Super-Resolution Microscopy Inside Live Cells Using Reversible Peptide-Protein Interactions"

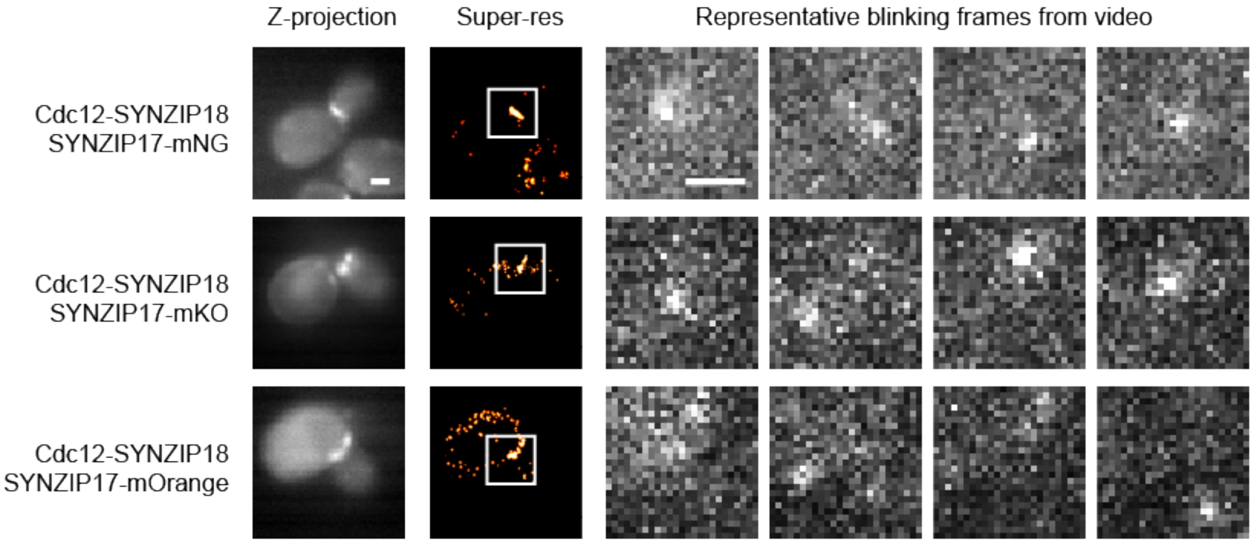


Figure S1. LIVE-PAINT can be performed with different FPs. From top to bottom, LIVE-PAINT was performed using the SYNZIP17-18 interaction pair and three different FPs: mNG, mKO, and mOrange, as indicated. (Left) Z-projections showing the average fluorescence signal for each video, calculated by integrating the average intensity over the entire video. (Middle) Super-resolution images for each video. The white box corresponds to the cropped region shown in the "representative blinking frames from video" section at right. (Right) Representative frames from the video, showing bright "blinks" in different locations. Number of localization events: mNG: 531; mKO: 280; mOrange: 154. Scale bars are 1 μm.

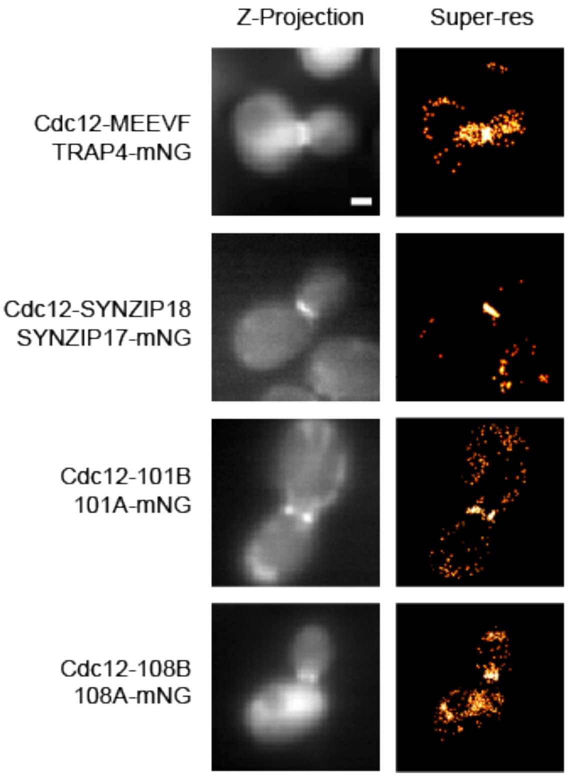


Figure S2. LIVE-PAINT can be performed with different peptide-protein interaction pairs. From top to bottom, the interaction pairs tested were TRAP4-MEEVF, SYNZIP17-SYNZIP18, 101A-101B, and 108A-108B, as indicated. (Left) Z-projections showing the average fluorescence signal for each video, calculated by integrating the average intensity over the entire video. (Right) Super-resolution images for each video. Number of localization events: TRAP4-MEEVF: 429; SYNZIP17-SYNZIP18: 398; 101A-101B: 582; 108A-108B: 803. Scale bars are 1 μm.

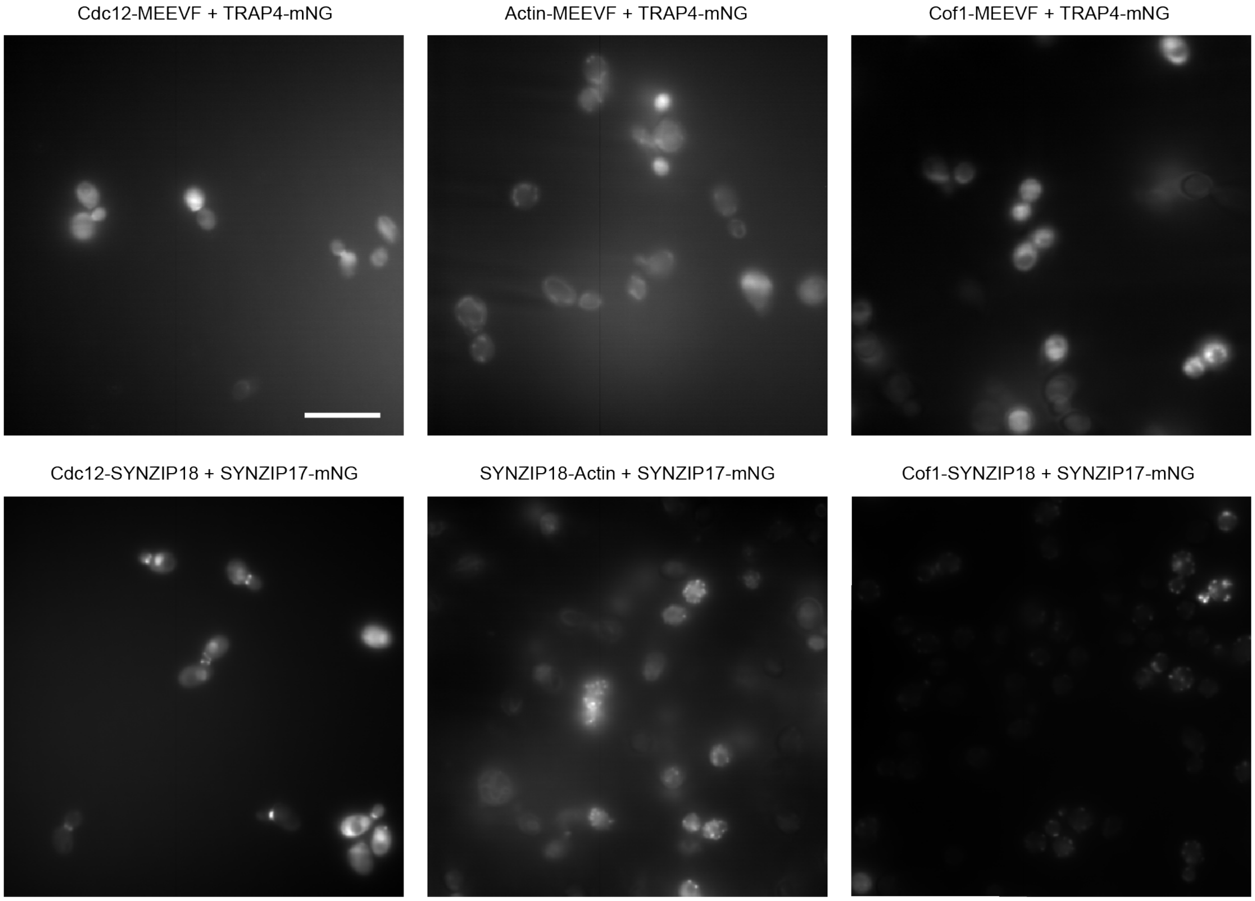


Figure S3. TRAP4-MEEVF and SYNZIP17-SYNZIP18 interactions are specific inside the cell. From left to right, maximum projection images of Cdc12, actin, and Cof1, which were tagged by either the TRAP4-MEEVF (top row) or SYNZIP17-SYNZIP18 (bottom row) interaction pair. Structures localized to the septum are seen when tagging Cdc12 and puncta around the edge of the cell are observed when tagging actin of Cof1. Scale bar is 10 μm.

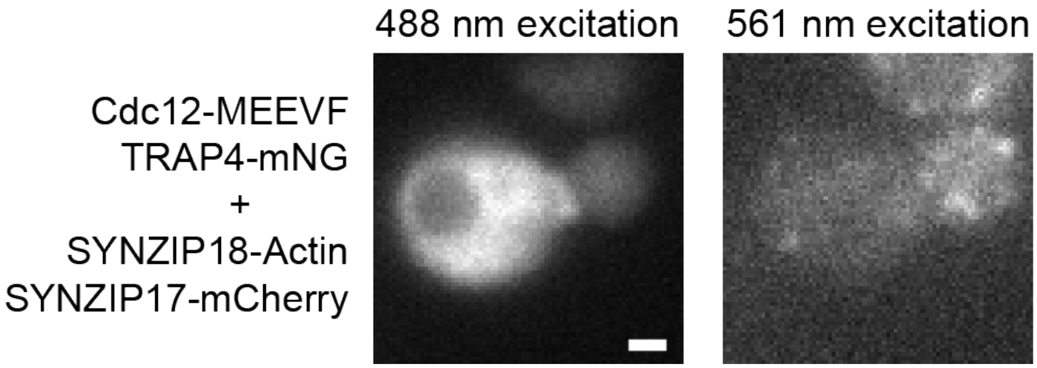


Figure S4. TRAP4-MEEVF and SYNZIP17-SYNZIP18 interaction pairs are orthogonal to one another and can be used with two different FPs for concurrent imaging. Fluorescence images of a cell expressing both Cdc12-MEEVF+TRAP4-mNG and SYNZIP18-Actin+SYNZIP17-mCherry. (Left) Cell imaged using a 488 nm excitation laser and green emission filter. Structure at yeast septum, corresponding to the location of Cdc12, is clearly visible. mNG fluorescence would be detected using these excitation and emission settings. (Right) Cell imaged using a 561 nm excitation laser. Distinctive structures around the edge of the cell, corresponding to the location of actin, are clearly visible. mCherry fluorescence would be detected using these excitation and emission settings. Images were collected using a 1 s exposure time. Scale bars are 1 μm.

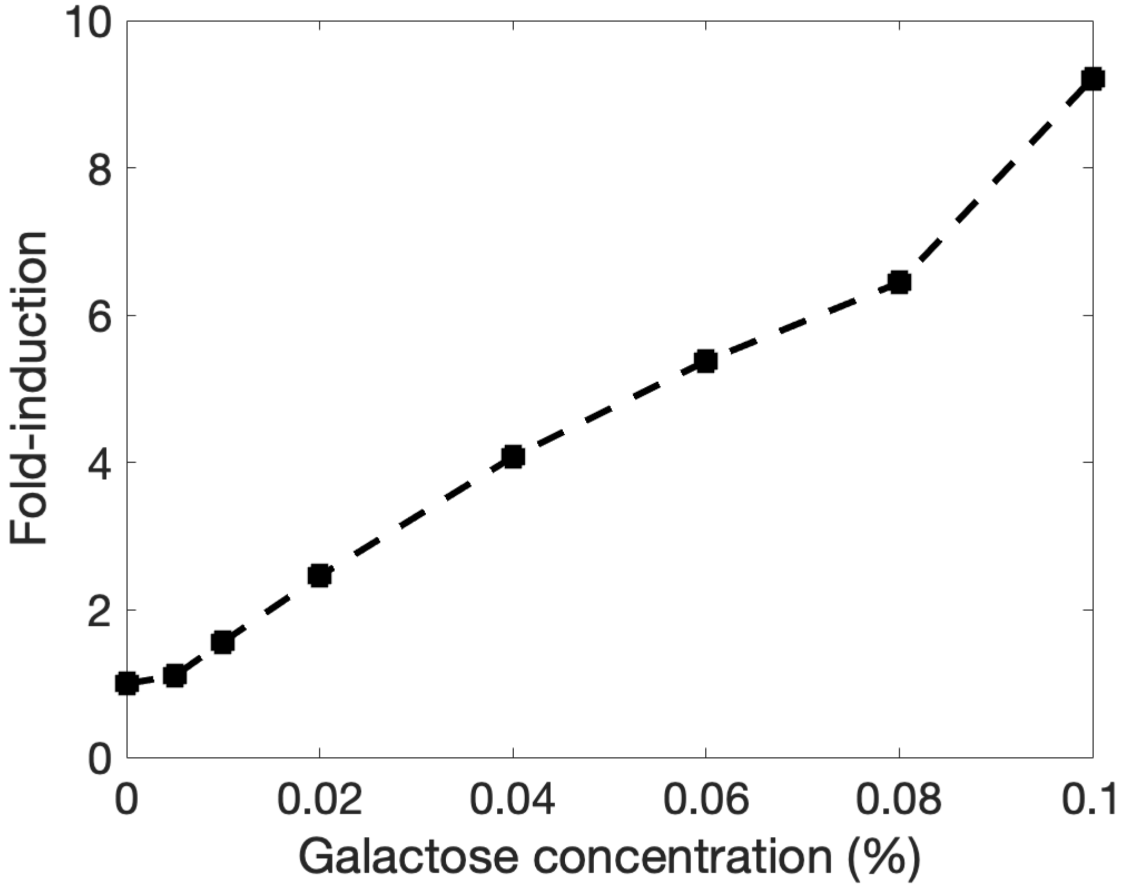


Figure S5. Expression of mNG under pGAL1 is linear with galactose concentration in gal2Δ background. SYNZIP17-mNG was expressed under pGAL1 and grown overnight in synthetic complete media supplemented with 1% raffinose and a variable amount of galactose. No glucose was added to the media, as glucose represses pGAL1. The expression of mNG was normalized first to the $OD_{600}$ of the culture, which was between 0.12 and 0.16. This fluorescence value was then normalized to the expression level at 0% galactose. At higher galactose concentrations (2%) we have seen fold-induction values of ~30.

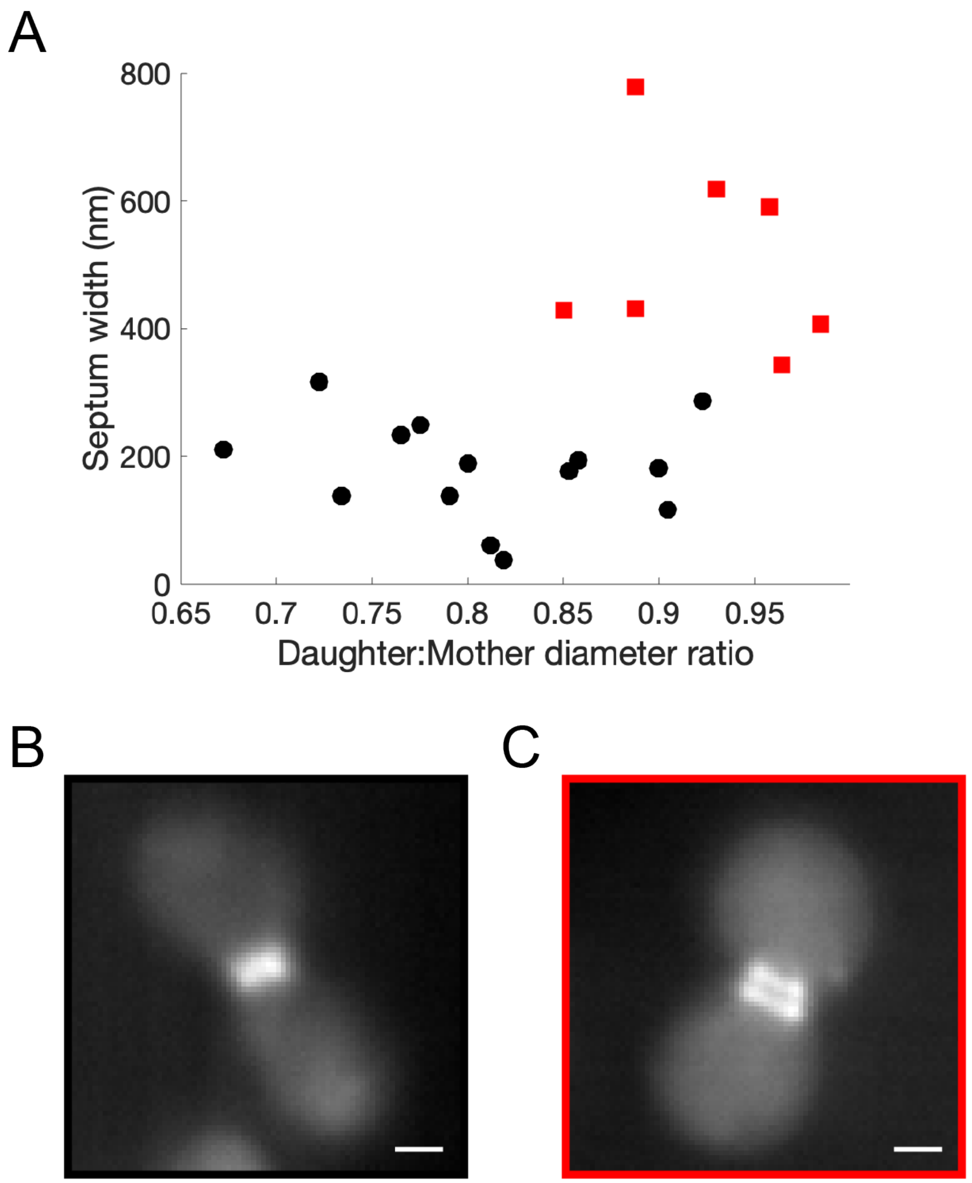


Figure S6. Yeast septum width increases with daughter:mother diameter ratio. (A) septum width plotted as a function of yeast daughter:mother diameter ratio, with single-ring septa plotted with black dots and double-ring septa plotted with red squares. See methods for how we determined septum width. Single-ring and double-ring septa were readily identifiable from fluorescence images of single cells. (B) and (C) show representative fluorescence images for single-ring and double-ring septa, respectively. Scale bars are 1 μm.

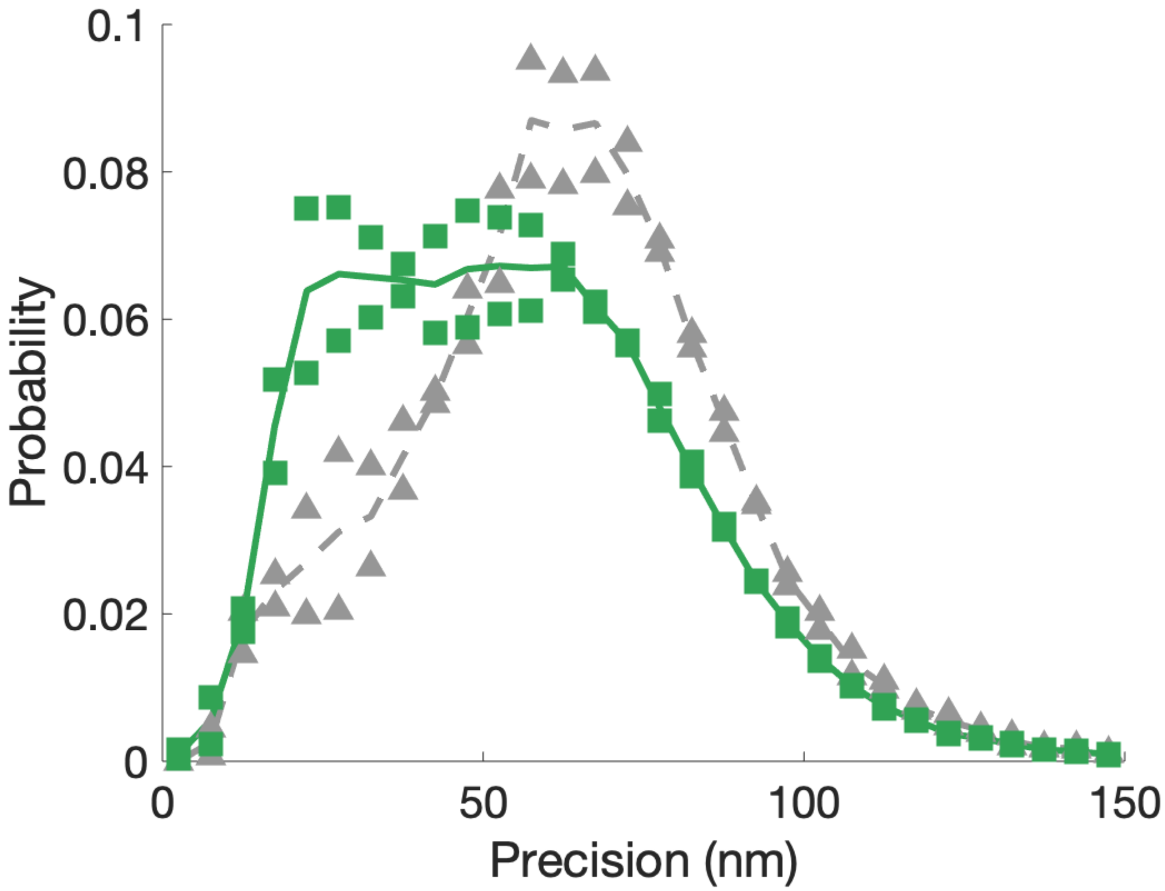


Figure S7. SYNZIP17-3xmNG shows improved localization precision compared with SYNZIP17-1xmNG. Full distribution of localization precision shown for Cdc12-SYNZIP18 + SYNZIP17-3xmNG (green squares and solid line) and Cdc12-SYNZIP18 + SYNZIP17-1xmNG (gray triangles and dashed line). Both experiments were performed by expressing the FP construct using 0.005% galactose in the yeast growth media. The curves show the average of two replicates for both the 3xmNG and 1xmNG constructs, while the data points for both replicates are given as spots.

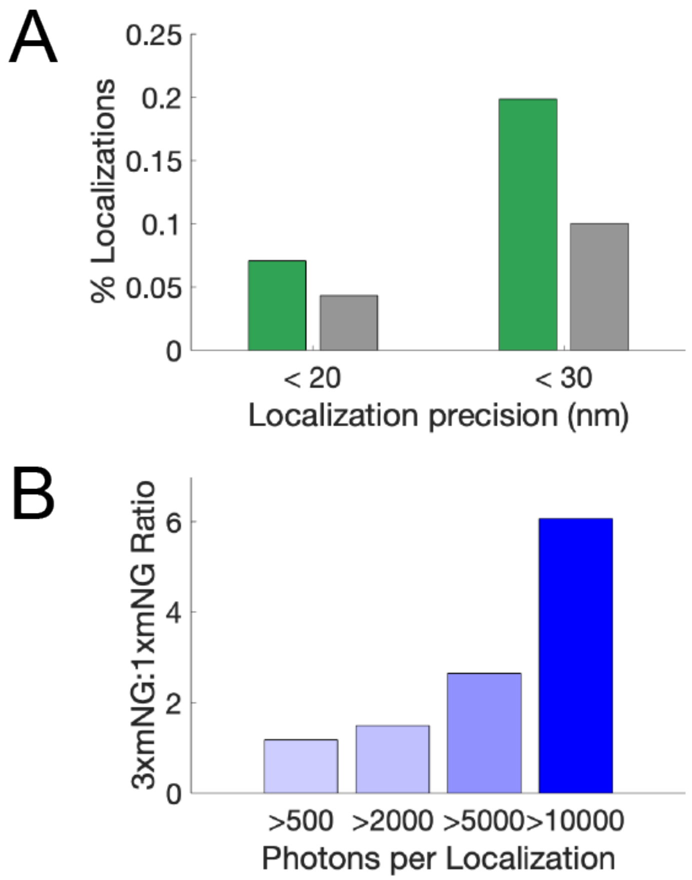


Figure S8. Three tandem copies of mNeonGreen (3xmNG) shows improved localization precision compared with a single copy of mNG fused to the cognate peptide binding protein (A) Percentage of localizations with precision values < 20 nm or < 30 nm. Green bars represent data for 3xmNG, and the gray bars represents data for 1xmNG. (B) LIVE-PAINT with 3xmNG gives higher numbers of localizations with a large number of photons than with mNG. The 3xmNG:mNG ratio of number of localizations is plotted for each 'photons per localization' bin. The darker the blue bar, the greater the enrichment in probability for 3xmNG compared with mNG in that bin.

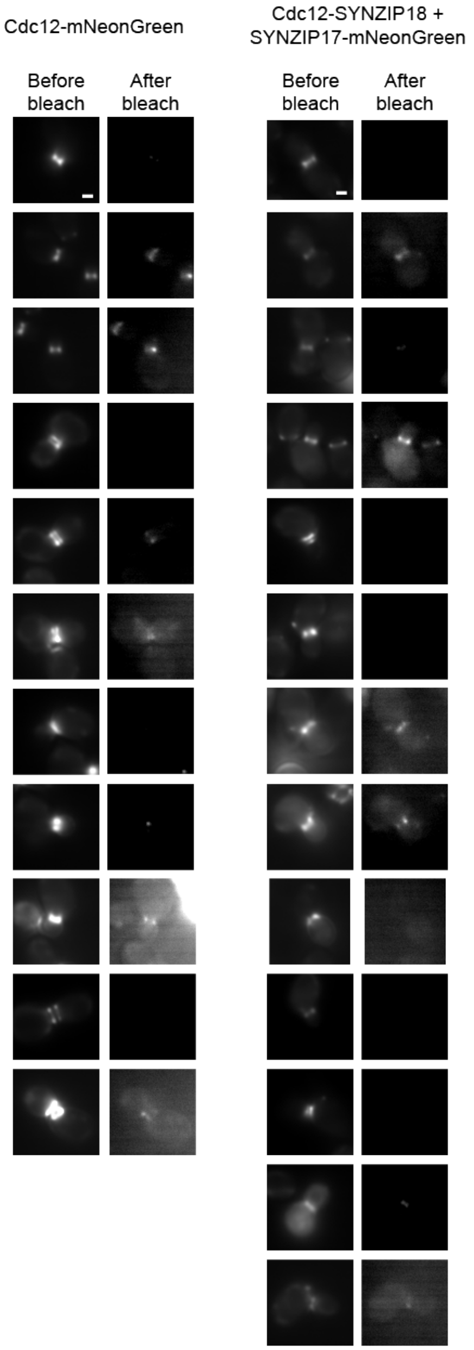


Figure S9. Maximum projection images generated from videos collected before and after bleaching, for all cells analyzed for the data shown in Figure 3. (Left) Maximum projection images are shown for cells expressing Cdc12-mNG. (Right) Maximum projection images are shown for cells expressing Cdc12-SYNZIP18 + SYNZIP17-mNG. Scale bar is 1 μm.

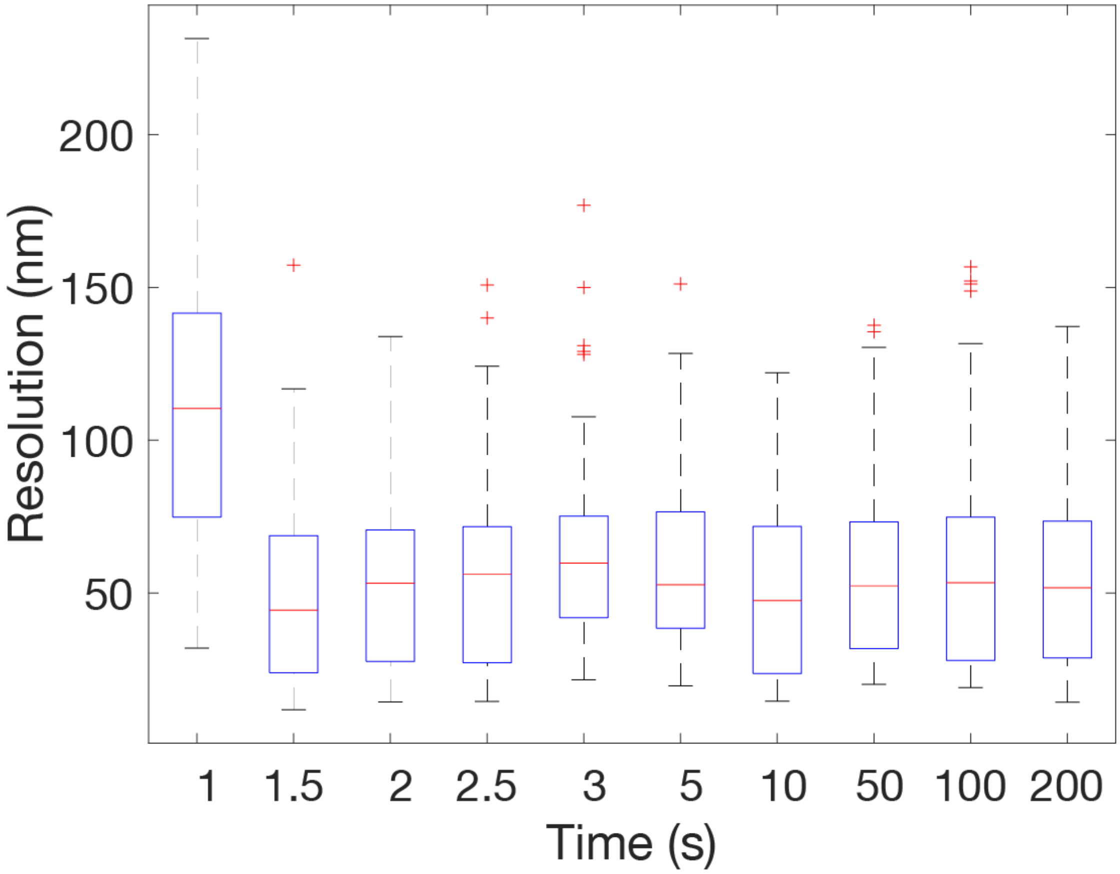


Figure S10. Image resolution for SYNZIP18-actin + SYNZIP17-mNG levels off at 50 nm after 1.5 s of data collection. Videos of different lengths indicated by the x-axis values were used to construct super-resolution images for the SYNZIP18-actin + SYNZIP17-mNG constructs. The resolution of the resulting images was measured using a Fourier Ring Correlation (FRC) method. This was performed 100 times for each image, resulting in the boxplot data shown above.

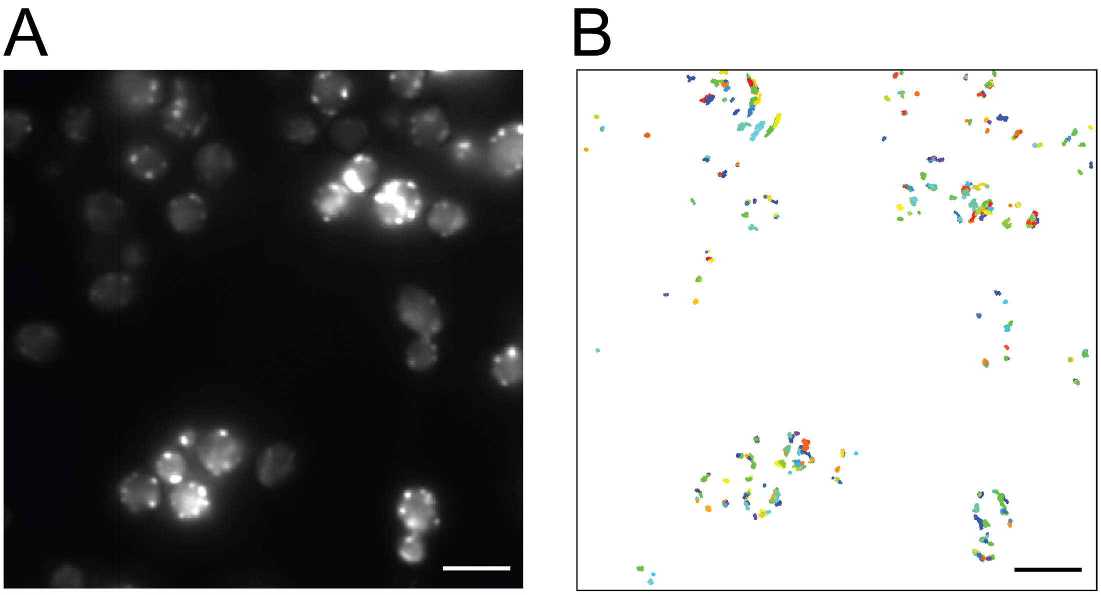


Figure S11. Tracking of cofilin in yeast cells. (A) Diffraction-limited image of the cofilin in the yeast cells. (B) Tracks from individual cofilin clusters. Scale bar is 5 μm.

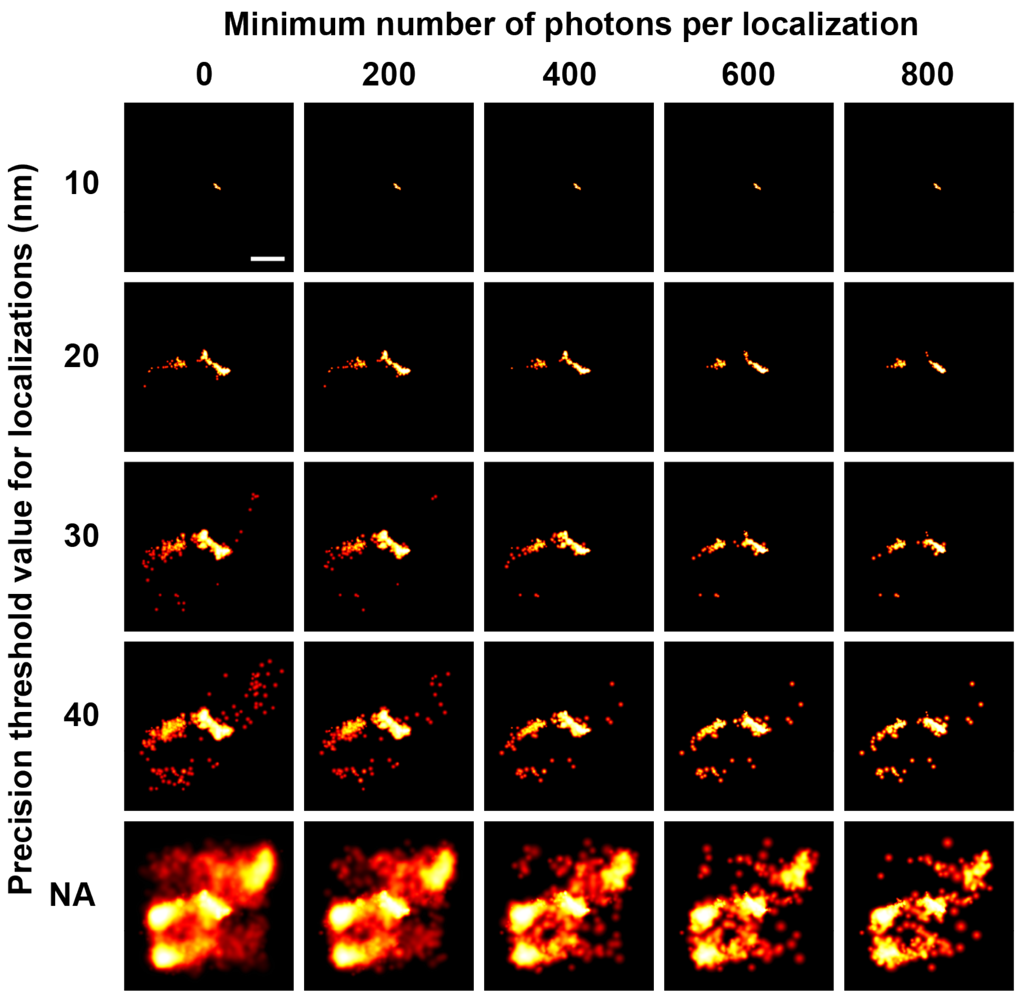


Figure S12. One stack of images analyzed using different thresholds for localization precision and minimum number of photons per localization. NA indicates no precision value specified. Scale bar is 1 μm.

**Supplementary Tables**

| Galactose concentration | TRAP % localisations in septum | TRAP Total number of localisations | SYNZIP % localisations in septum | SYNZIP Total number of localisations |
|---|---|---|---|---|
| 0 % | 14.71 | 272 | 94.12 | 153 |
| 0.005 % | 44.54 | 348 | 97.99 | 398 |
| 0.02% | 37.76 | 429 | 42.69 | 878 |
| 0.1% | 22.78 | 1203 | 18.51 | 1313 |

Table S1. Quantification of the total number of super-resolution localisations and the percentage of these localisations in the septum for images shown in Figure 2.

Table S2. Yeast strains used in this study

| Strain | Parent | Genotype | Reference |
|---|---|---|---|
| BY4741 | - | MATa his3Δ1 leu2Δ0 met15Δ0 ura3Δ0 | Marsden Lab |
| CDC12-MEEVF | BY4741 | MATa his3Δ1 leu2Δ0 met15Δ0 ura3Δ0 CDC12-MEEVF::KanMX6 | This study |
| CDC12-SYNZIP18 | BY4741 | MATa his3Δ1 leu2Δ0 met15Δ0 ura3Δ0 CDC12-SYNZIP18::KanMX6 | This study |
| CDC12-CCBN3,5 | BY4741 | MATa his3Δ1 leu2Δ0 met15Δ0 ura3Δ0 CDC12-CCBN3,5::KanMX6 | This study |
| pGAL1 TRAP4-mEOS | BY4741 | MATa his3Δ1 leu2Δ0 met15Δ0 ura3Δ0 gal2Δ::His3MX6 pGAL1 TRAP4-mEOS | This study |
| pGAL1 TRAP4-mNG | BY4741 | MATa his3Δ1 leu2Δ0 met15Δ0 ura3Δ0 gal2Δ::His3MX6 pGAL1 TRAP4-mNG | This study |
| pGAL1 SYNZIP-mEOS | BY4741 | MATa his3Δ1 leu2Δ0 met15Δ0 ura3Δ0 gal2Δ::His3MX6 pGAL1 SYNZIP17-mEOS | This study |
| pGAL1 SYNZIP-mNG | BY4741 | MATa his3Δ1 leu2Δ0 met15Δ0 ura3Δ0 gal2Δ::His3MX6 pGAL1 SYNZIP17-mNG | This study |
| CDC12-MEEVF + pGAL1 TRAP4-mEOS | BY4741 | MATa his3Δ1 leu2Δ0 met15Δ0 ura3Δ0 gal2Δ::His3MX6 pGAL1 TRAP4-mEOS CDC12-MEEVF::KanMX6 | This study |

| | | | |
|---|---|---|---|
| CDC12-MEEVF + pGAL1 TRAP4-mNG | BY4741 | MATa his3Δ1 leu2Δ0 met15Δ0 ura3Δ0 gal2Δ::His3MX6 pGAL1 TRAP4-mNG CDC12-MEEVF::KanMX6 | This study |
| CDC12-SYNZIP18 + pGAL1 SYNZIP17-mEOS | BY4741 | MATa his3Δ1 leu2Δ0 met15Δ0 ura3Δ0 gal2Δ::His3MX6 pGAL1 SYNZIP17-mEOS CDC12-SYNZIP18::KanMX6 | This study |
| CDC12-SYNZIP18 + pGAL1 SYNZIP17-mNG | BY4741 | MATa his3Δ1 leu2Δ0 met15Δ0 ura3Δ0 gal2Δ::His3MX6 pGAL1 SYNZIP17-mNG CDC12-SYNZIP18::KanMX6 | This study |
| CDC12-SYNZIP18 + pGAL1 SYNZIP17-3xmNG | BY4741 | MATa his3Δ1 leu2Δ0 met15Δ0 ura3Δ0 gal2Δ::His3MX6 pGAL1 SYNZIP17-3xmNG CDC12-SYNZIP18::KanMX6 | This study |
| CDC12-CCBN3,5 + pGAL1 CCAN3,5-mEOS | BY4741 | MATa his3Δ1 leu2Δ0 met15Δ0 ura3Δ0 gal2Δ::His3MX6 pGAL1 CCAN3,5-mEOS CDC12-CCBN3,5::KanMX6 | This study |
| COF1-MEEVF + pGAL1 TRAP4-mNG | BY4741 | MATa his3Δ1 leu2Δ0 met15Δ0 ura3Δ0 gal2Δ::His3MX6 pGAL1 TRAP4-mNG COF1-MEEVF::KanMX6 | This study |
| COF1-SYNZIP18 + pGAL1 SYNZIP17-mNG | BY4741 | MATa his3Δ1 leu2Δ0 met15Δ0 ura3Δ0 gal2Δ::His3MX6 pGAL1 SYNZIP17-mNG COF1-SYNZIP18::KanMX6 | This study |

Table S3. Sequencing Primers

| Label | Name | Sequence | Purpose |
|---|---|---|---|
| S1 | CDC12_CT_F | GAGGGTCACGAGAACACC | Check C-terminus of CDC12 |
| S2 | CDC12_CT_R | CAGTTACTTCTGCTGGTTCC | Check C-terminus of CDC12 |
| S3 | GAL2_Seq_F | CTAATCCAAGGAGGTTTAC | Check GAL2 locus |
| S4 | GAL2_Seq_R | TAAGAGAGATGATGGAGC | Check GAL2 locus |
| S5 | SP6_Seq_F | ATTTAGGTGACACTATAG | Sequence pFA6a-HIS3MX6 and pFA6a-KANMX6 plasmids |
| S6 | T7_Seq_F | TAATACGACTCACTATAGGG | Sequence pFA6a-HIS3MX6 and pFA6a-KANMX6 plasmids |
| S7 | HIS_SEQ_F | CGTTAGAACGCGGCTAC | Sequence pFA6a-HIS3MX6 plasmids |
| S8 | GAL2_SEQ_F2 | GCTGCAGAAGGCACATCTA | Check GAL2 locus |
| S9 | GAL2_SEQ_R2 | CCCAGAGATAAGTCTGGTGATG | Check GAL2 locus |
| S10 | pCUP1_seq_F | CATATAGAAGTCATCGACTAGT | Check pCu415CUP1 plasmid |
| S11 | pCUP1_seq_R | GACGGTATCGATAAGCTT | Check pCu415CUP1 plasmid |
| S12 | COF1_seq_F | CCTTAAACGGTGTCTCTACC | Check C-terminus of COF1 |

| | | | |
|---|---|---|---|
| S13 | COF1_seq_R | GGTGTACGGGACCTTAAATC | Check C-terminus of COF1 |

Table S4. Primers for amplifying plasmid backbone

| Label | Name | Sequence | Purpose |
|---|---|---|---|
| C1 | p6k_ath1_F | TTGCAAACCAGAGCCTG | Amplify pFA6a-KANMX6 backbone to replace MEEVF with another peptide |
| C2 | p6k_ath1_R | TGATGAGTCATGTAATTAGTTATGT | Amplify pFA6a-KANMX6 backbone to replace MEEVF with another peptide |
| C3 | p6h_ath2_F | GAATCCGGGGTTTTTTCT | Amplify pFA6a-HIS3MX6 backbone to replace TRAP4 with another protein |
| C4 | p6h_ath2_R | CTGCAGATGAGTGCGATTA | Amplify pFA6a-HIS3MX6 backbone to replace TRAP4 with another protein |
| C5 | p6h_ath3_t4_F | TTGCTCCTTCAGGATTTTCT | Amplify pFA6a-HIS3MX6 backbone to replace TRAP4 mEOS with another FP (e.g. mNG). TRAP4 is left to make a TRAP4-FP fusion. |
| C6 | p6h_ath3_sz_F | CTTGTAAGCTTCAATTTCCTTTCTCAAGT | Amplify pFA6a-HIS3MX6 backbone to replace SYNZIP17 mEOS with another FP (e.g. mNG). |

| | | | |
|---|---|---|---|
| | | | SYNZIP17 is left to make a SYNZIP17-FP fusion. |
| C7 | p6h_ath3_R | TGATAAGTCATGTAATTAGTTATGTC | Amplify pFA6a-HIS3MX6 backbone to replace TRAP4 or SYNZIP mEOS with another FP (e.g. mNG). TRAP4 or SYNZIP17 is left to make a TRAP4/SYNZIP17-FP fusion. |
| C8 | pCUP1_ATH_F | AAGCTTATCGATACCGTC | Amplify pCu415CUP1 backbone to replace product under expression on pCUP1 |
| C9 | pCUP1_ATH_R | ACTAGTCGATGACTTCTATATG | Amplify pCu415CUP1 backbone to replace product under expression on pCUP1 |
| C10 | p6h_ath3_mNG_R | ATAACATGGCCTCTCTCC | Amplify pFA6a-HIS3MX6 backbone to replace TRAP4-mNG with LifeAct-mNG (use with C3) |

Table S5. Integration primers

| Label | Name | Sequence | Purpose |
|---|---|---|---|
| I1 | p6h_int_F | CTAATCCAAGG AGGTTTACGGA CCAGGGGAAC TTTCCAGATTC AGAAGCTTCGT ACGCTGCA | Amplify the entire cassette from pFA6a-HIS3MX6 (e.g. TRAP4-mEOS under GAL1, plus HIS3 marker) for transformation into yeast |
| I2 | p6h_int_R | CATGAAAAATT AAGAGAGATGA TGGAGCGTCTC ACTTCAAACGC AGGCGTTAGTA TCGAATCG | Amplify the entire cassette from pFA6a-HIS3MX6 (e.g. TRAP4-mEOS under GAL1, plus HIS3 marker) for transformation into yeast |
| I3 | p6k_int_F | GAAGAGCAGG TCAAAAGCTTG CAAGTAAAAAA ATCCCATTTAA AAGGTGGATCA GGCTCTGG | Amplify the entire cassette from pFA6a-KANMX6 (e.g. GS-MEEVF, plus KANMX6 marker) for transformation into yeast |
| I4 | p6k_int_R | AGGCGTTGAAA TTGACGAGACA AAGAGGAAGA | Amplify the entire cassette from pFA6a-KANMX6 (e.g. GS-MEEVF, plus KANMX6 marker) for transformation into yeast |

| | | | |
|---|---|---|---|
| | | CATTAATTAAT CATTAGAAAAA CTCATCGAGCA TC | |
| I5 | p6kcof1intF | TACGATTCTGT TTTGGAAAGAG TCAGCAGAGG CGCTGGTTCTC ATGGTGGATCA GGCTCTGG | Amplify the entire cassette from pFA6a-KANMX6 (e.g. GS-MEEVF, plus KANMX6 marker) for transformation into yeast at Cof1 locus |
| I6 | p6kcof1intR | TTTCATTTTTCT TGAAGATTGTT GTCATTTGTGA AATCATTTACC ATTAGAAAAAC TCATCGAGCAT C | Amplify the entire cassette from pFA6a-KANMX6 (e.g. GS-MEEVF, plus KANMX6 marker) for transformation into yeast at Cof1 locus |
